# Quantitative Action Spectroscopy Reveals ARPE-19 Sensitivity to Long-Wave Ultraviolet Radiation at 350 nm and 380 nm

**DOI:** 10.1101/2021.12.09.471589

**Authors:** Graham Anderson, Andrew McLeod, Pierre Bagnaninchi, Baljean Dhillon

## Abstract

The role of ultraviolet radiation (UVR) exposure in the pathology of age-related macular degeneration (AMD) has been debated for decades with epidemiological evidence failing to find a clear consensus for or against it playing a role. A key reason for this is a lack of foundational research into the response of living retinal tissue to UVR in regard to AMD-specific parameters of tissue function. We therefore explored the response of cultured retinal pigmented epithelium (RPE), the loss of which heralds advanced AMD, to specific wavelengths of UVR across the UV-B and UV-A bands found in natural sunlight.

Using a bespoke *in vitro* UVR exposure apparatus coupled with bandpass filters we exposed the immortalised RPE cell line, ARPE-19, to 10nm bands of UVR between 290 and 405nm. Physical cell dynamics were assessed during exposure in cells cultured upon specialist electrode culture plates which allow for continuous, non-invasive electrostatic interrogation of key cell parameters during exposure such as monolayer coverage and tight-junction integrity. UVR exposures were also utilised to quantify wavelength-specific effects using a rapid cell viability assay and a phenotypic profiling assay which was leveraged to simultaneously quantify intracellular reactive oxygen species (ROS), nuclear morphology, mitochondrial stress, epithelial integrity and cell viability as part of a phenotypic profiling approach to quantifying the effects of UVR.

Electrical impedance assessment revealed unforeseen detrimental effects of UV-A, beginning at 350nm, alongside previously demonstrated UV-B impacts. Cell viability analysis also highlighted increased effects at 350nm as well as 380nm. Effects at 350nm were further substantiated by high content image analysis which highlighted increased mitochondrial dysfunction and oxidative stress.

We conclude that ARPE-19 cells exhibit a previously uncharacterised sensitivity to UV-A radiation, specifically at 350nm and somewhat less at 380nm. If upheld *in vivo*, such sensitivity will have impacts upon geoepidemiological risk scoring of AMD.

## Introduction

The question of if, and if so to what degree, solar radiation exposure is involved in the pathogenesis of age-related macular degeneration (AMD) has long been the subject of medical contemplation and investigation [1] with academic interest in the subject widely piqued in the latter half of the 20^th^ century which ushered in high-altitude flight and atomic ordinance capable of significant retinal injury [2]. However, investigations of the time relied upon a small number of large animals and semi-quantitative means of assessing UVR damage both of which negatively impact upon the statistical power and sensitivity of the observations in describing sub-acute perturbations[3].

Consequently, contemporary estimates regarding solar photosensitivity of the retina generally favour UVR wavelengths known to be poorly transmitted to the retinal surface[4,5], thus leading to the conclusion that ‘harmful UVR’ does not reach the retina, despite evidence highlighting paediatric transmission of UV-A [5]. Moreover, the role of lifetime sun exposure in retinal degeneration has been explored by Schick *et al*. [10] who used questionnaires to determine that sun exposure within the paediatric and occupationally active years of a person’s life are correlated with the onset of AMD in later years. As a result, recent *in vitro* investigations utilising fully quantitative methods of characterising UVR exposure effects tend to under sample the UV-A band in favour of high-energy visible (HEV; 400-470nm) and UV-B wavelengths, and in some cases highly biologically effective UV-C wavelengths which do not reach the Earth’s surface within sunlight [6–9].

Ultimately, the quantitative evidence-base regarding solar UVR (280-400nm) driven effects within the retina, in terms congruent with modern oxidative-stress theories of ageing and disease, remains incomplete. This impedes the accurate geographic modelling of ophthalmologically harmful UVR reaching the Earth’s surface which, in turn, hampers ongoing geo-epidemiological risk modelling of AMD and the shepherding of resources to meet future ophthalmic needs.

In the present study, we sought to characterise the influence of solar UVR upon the cells centrally implicated in the pathogenesis of AMD, the retinal pigmented epithelium (RPE), using fully quantitative methods to determine cell viability, oxidative stress burden and tight-junction integrity with a high degree of spectral resolution to create response spectra which can be used in the modelling of ophthalmic risk within the scope of a changing climate and large scale demographic processions as well as to infer putative molecules of interest in UVR damage to the RPE.

## Results

### Full Spectrum Viability Assay Suggests Unique UV-A Effects

Initial investigations - coupling a bespoke *in vitro* exposure apparatus to irradiate cells and a rapid cell viability agent to quantify wavelength-specific effects - highlighted the clear distinction in photo-damage between the UV-B and UV-A bands (*Figure 3*; panel a) with the UV-B wavelengths exhibiting toxic efficiencies several orders of magnitude greater than the UV-A or visible bands (*Figure 3*; panel b)).

**Figure 1.**
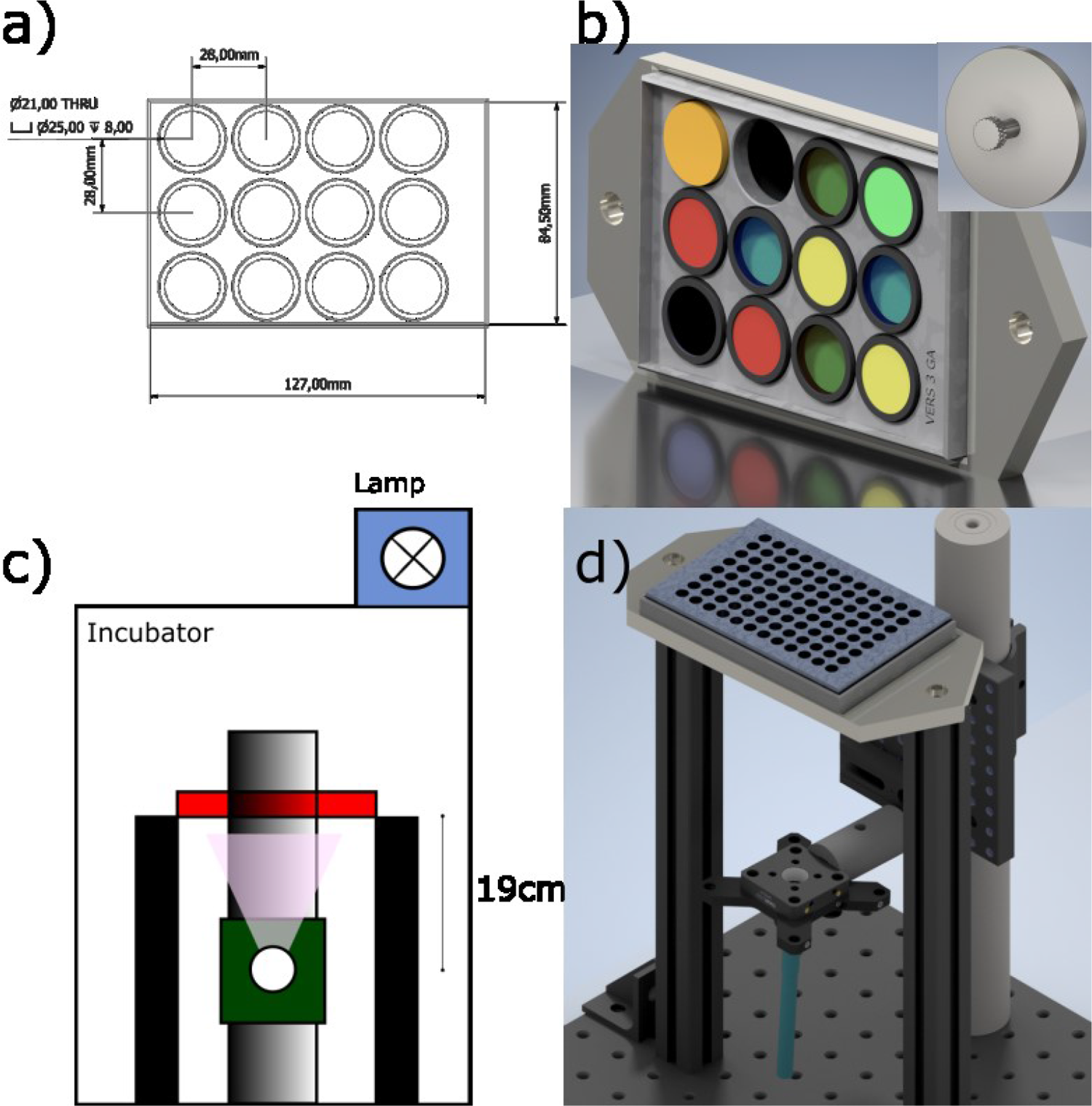
Outline of Exposure Apparatus. a) Technical drawing of bandpass filter holder used during exposure. b) 3D render of filter holder with filters inserted. Inset 3D render of ‘blank’ filter which provided a negative control. c) Schematic of overall design highlighting the position of the culture plate (red box) in relation to the light source (white circle). d) Computer generated mock-up of full optomechanical apparatus with culture plate in place.

**Figure 2.**
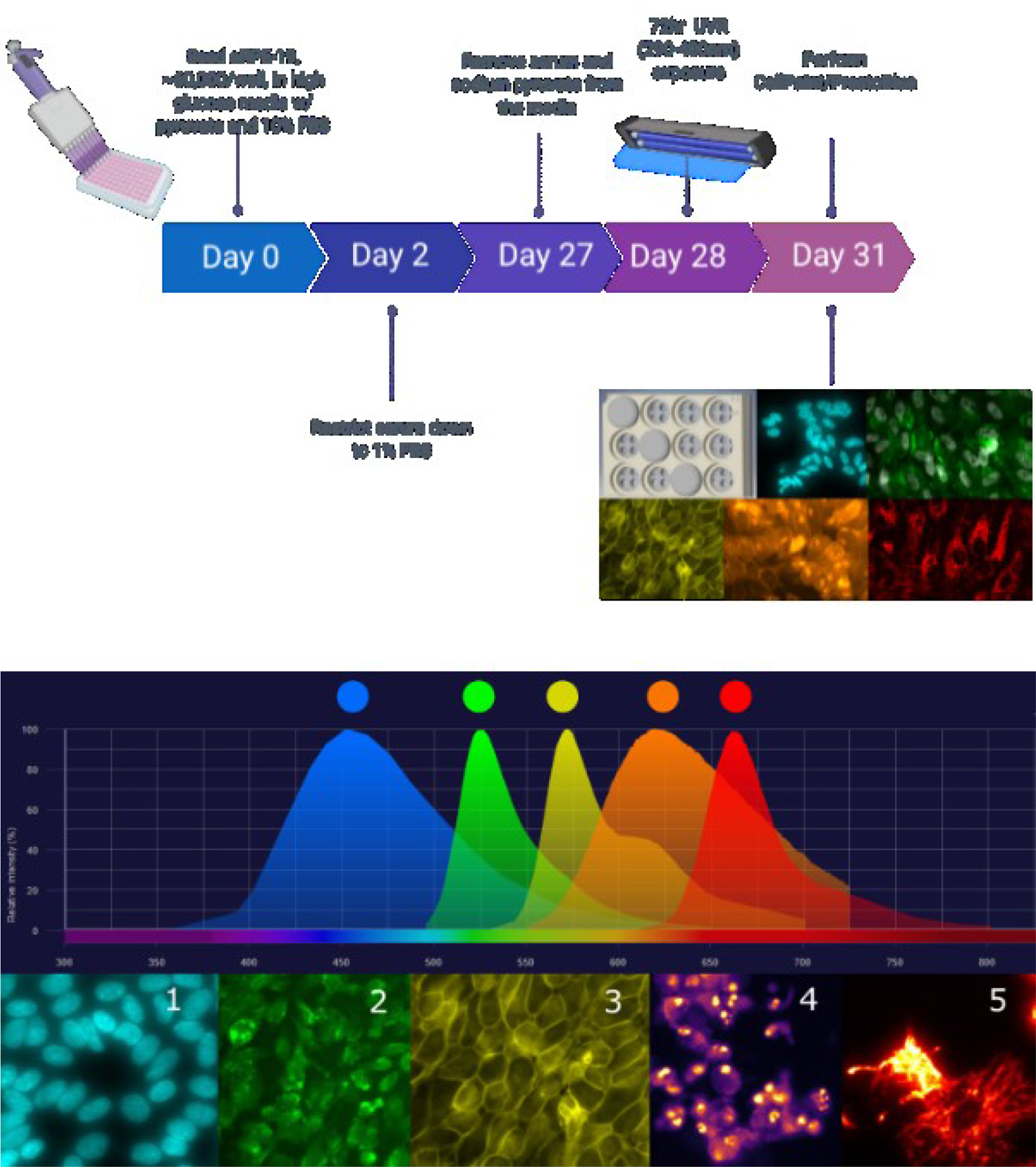
Top - The prototypical workflow of the current project. Mid - The emission spectra of the fluorescent probes used in the current study. Bottom - representative images in ascending order of excitation wavelength. 1 = Hoechst 33342; 2 = carboxy-H2DCFDA; 3 = CellMask™ Orange; 4 = Propidium Iodide; 5 = MitoTracker™ Deep Red FM.

**Figure 3.**
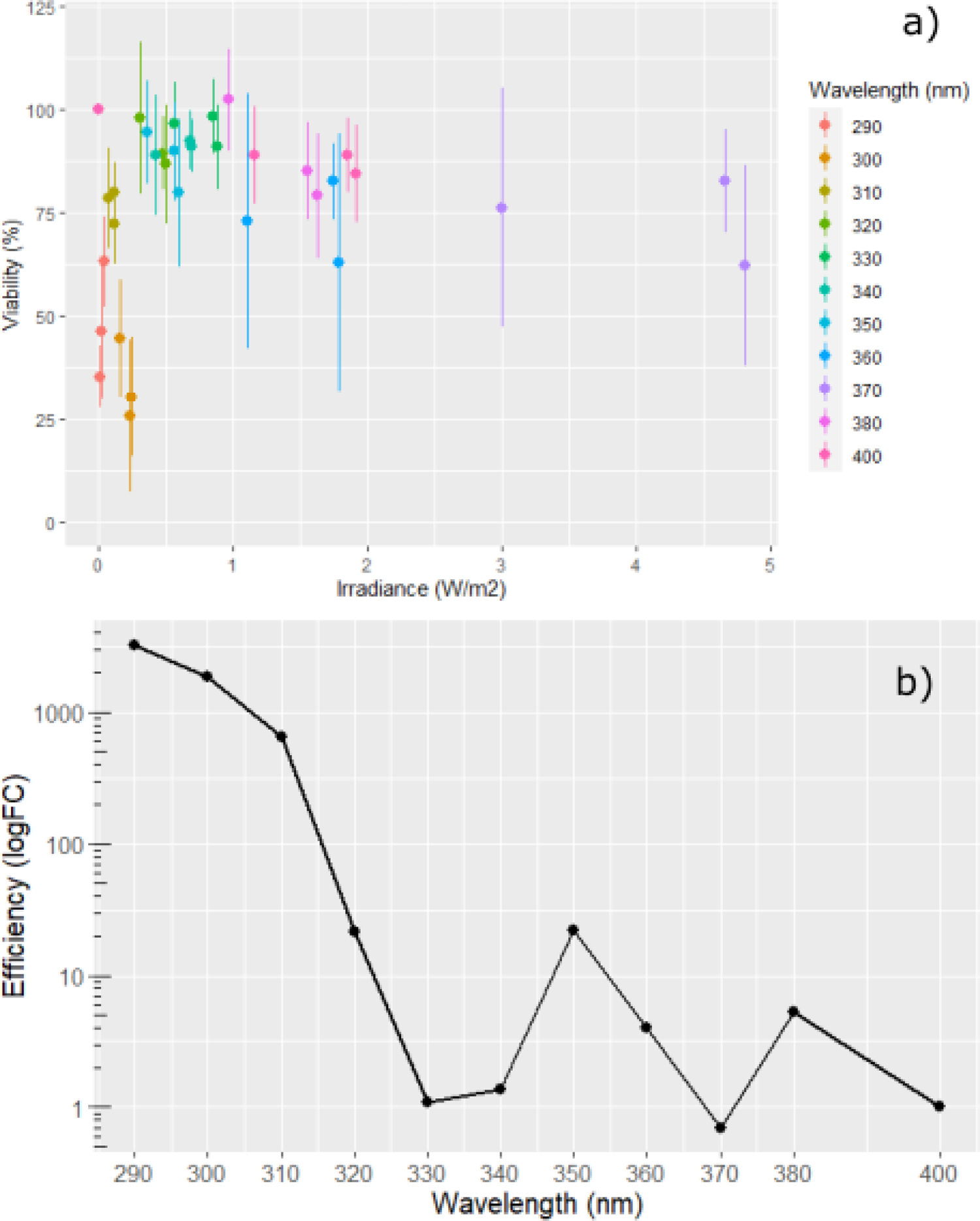
Average cell viability, assessed via Prestoblue™, as a function of irradiance and wavelength. b) The resultant response spectrum based on the slope coefficients of the viability data. N = 3, Error bars = 1 s.d.

While UV-A radiation was well tolerated by ARPE-19, with a mean decrease of cell viability over the radiation band of 20% of control, response spectra analysis revealed distinct peaks in effects at 350 nm and 380 nm for which a 1.6-fold increase in intensity resulted in a 10% and 30% decrease in cell viability, presenting an approximately 5 and 10-fold increase respectively from effects at 405 nm (*Figure 3*; panel b).

### ECIS Action Spectra Highlights Response to UV-A Band

After four weeks in culture, immediately prior to UVR exposure, the media the cells were maintained in was replaced with custom media free from antioxidants and photosensitisers. Following this, the electrode plate the cells were cultured upon was docked in the 96-well ECIS station, the lid removed, and the plate exposed to the UVR source continuously for 68-72 h (*Figure 4*; panel a)).

**Figure 4.**
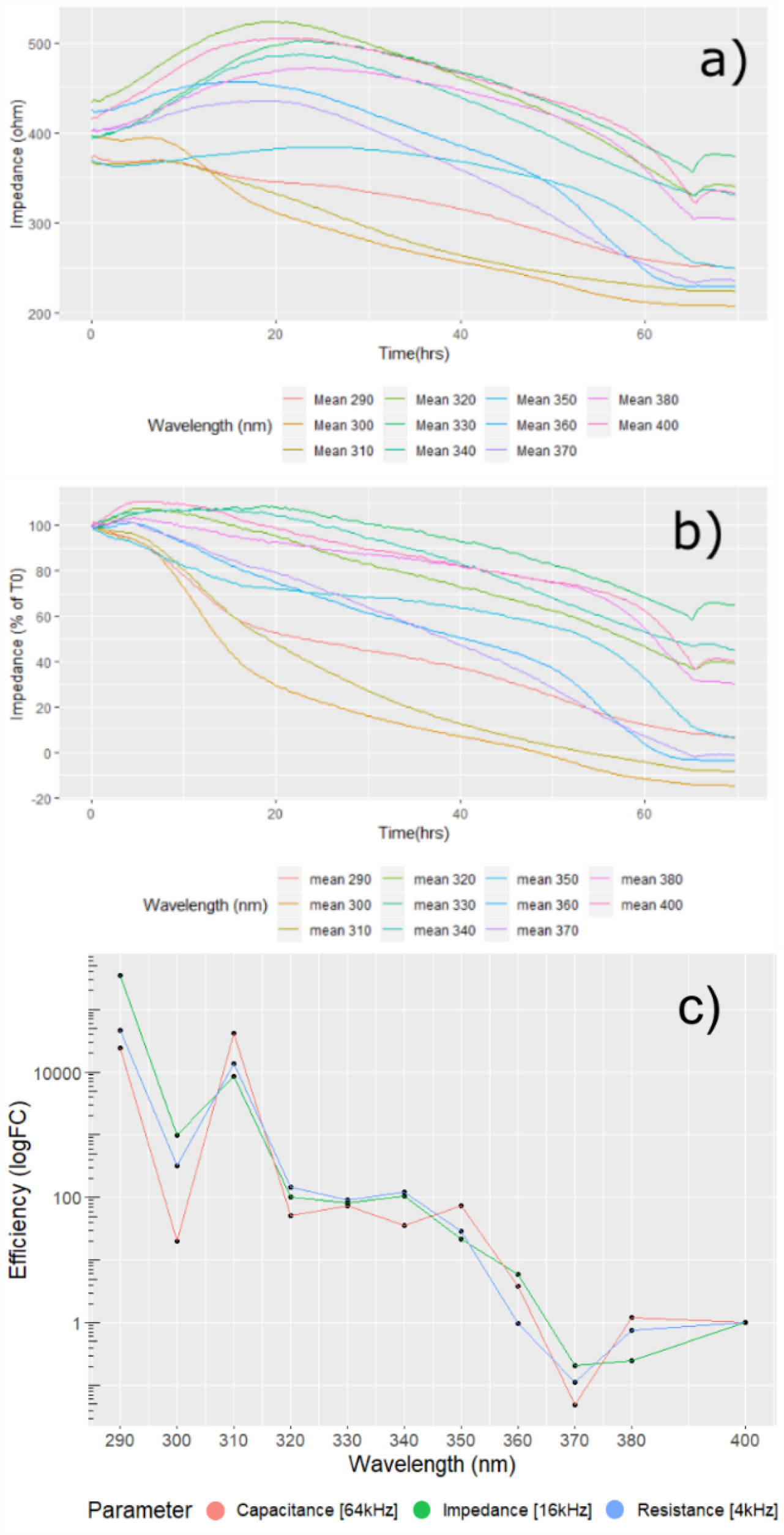
ECIS Data Processing. a) Raw impedance data [16kHz] averaged for each wavelength. Note the rapid rise in signal at 68hrs highlighting the time at which the lamp was doused. b) Impedance [16kHz] data following normalisation to time-zero (T0) and scaling between positive and negative controls. c) Resultant action spectra, normalised to 400nm, indicating the relative efficiency of each wavelength in reducing each electrostatic parameters to 60% of its initial value. N = 3.

Within the first 24 hours, the majority of wavelengths exhibited a rise in impedance typical of cell responses to changes in ionic balance and CO_2_ following a media change. The exception to this lies in the cells which were irradiated with UV-B radiation, where the inflection in electrostatic response was observed within the first 24 hours (*Figure 2*; panel b)).

Of note when considering the cellular impedance responses to particular wavelengths is the biphasic relationship which emerged over time. This was particularly noticeable in the response to 350 nm UVR which showed a rapid decrease in the first 6 hours of exposure followed by an extended plateau until around 45 hours where cellular impedance displayed a precipitous decline. This biphasic response suggests two separate events taking place, most likely an initial break down of cellular tight junctions followed by the eventual dissolution of cellular filopodia leading to detachment of the cells from their substrate.

Following normalisation to positive (media-only) and negative (No UVR) controls, action spectra were constructed based on the effective dose required to decrease the electrostatic parameter of interest (impedance [Z], resistance [R] and capacitance [C]) to 60% of their respective time-zero (t_0_) value (*Figure 4*; panel c)). The action spectra revealed that longwave UV-A, UV-A1, was well tolerated with an apparent trough in toxicity at 370 nm suggesting that, in regard to tight junctional breakdown, it is at least as toxic as high-energy visible (HEV) light but could be slightly beneficial to epithelial barrier function. A plateau of toxicity, around 100-fold HEV toxicity, was observed within the shortwave UV-A, UV-A2, band beginning at 350nm and extending to 320nm followed by a typical rise in toxicity within the UV-B band reaching 10,000-fold of HEV.

### Full Depth Phenotypic Profiling Recapitulates 350 nm Peak

Initially, variables of interest were chosen according to biological understanding of the processes of UV-B and UV-A pathology. However, following this, an open-ended data-driven approach was employed where all 380 imaging variables were modelled and included in the resultant response spectra.

When considering the overall cell response, that is, the average of all the image parameters for each stain, the resulting response spectra correlates well with the response spectra produced via the PrestoBlue™ (Invitrogen™, MA, USA) and nuclei count assays (*Figure 5*; *plot a)*. This suggests that nuclei staining, and the resulting intensity and morphological parameters, provide the most sensitive metric for overall UVR response. Dividing the data by the primary imaging parameters determined by probe: nucleus, mitochondria, reactive oxygen species (ROS), propidium iodide/Cell mask (PI/Cm) it becomes possible to discern which features diverge most from the overall modelled response at each wavelength (*Figure 5*; *plots b & c*).

**Figure 5.**
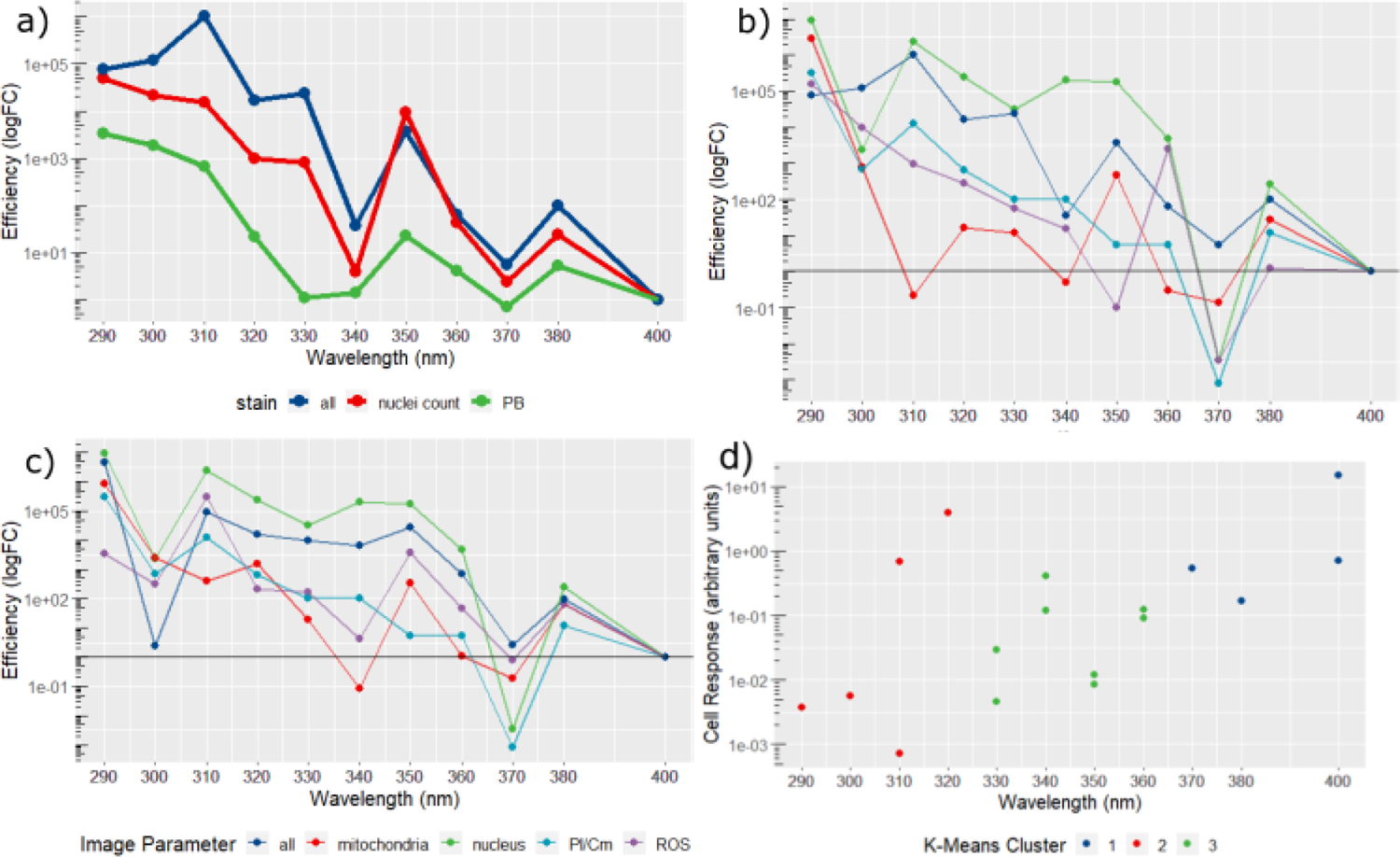
High Content Image Analysis. a) Response spectrum created using all available imaging parameters, the 350nm and 380nm peaks observed in previous data can be seen. b) Response spectra from imaging data broken down into major families: All, Mitochondria, Nucleus, Propidium Iodide/Cell Mask (PI/Cm), Reactive Oxygen Species (ROS). c) Response spectra created using imaging data produced by the ‘spot’ filtering kernel which successfully recapitulates the 350nm peak in every compartment except PI/Cm. d) Mapping clusters (k3) onto overall cell response differentiates three main groups, one comprising the UV-B band (290-320nm) and two within the UV-A band (330-360nm; 370-400nm) possibly indicating discrete mechanisms of action. N = 3

**Figure 6.**
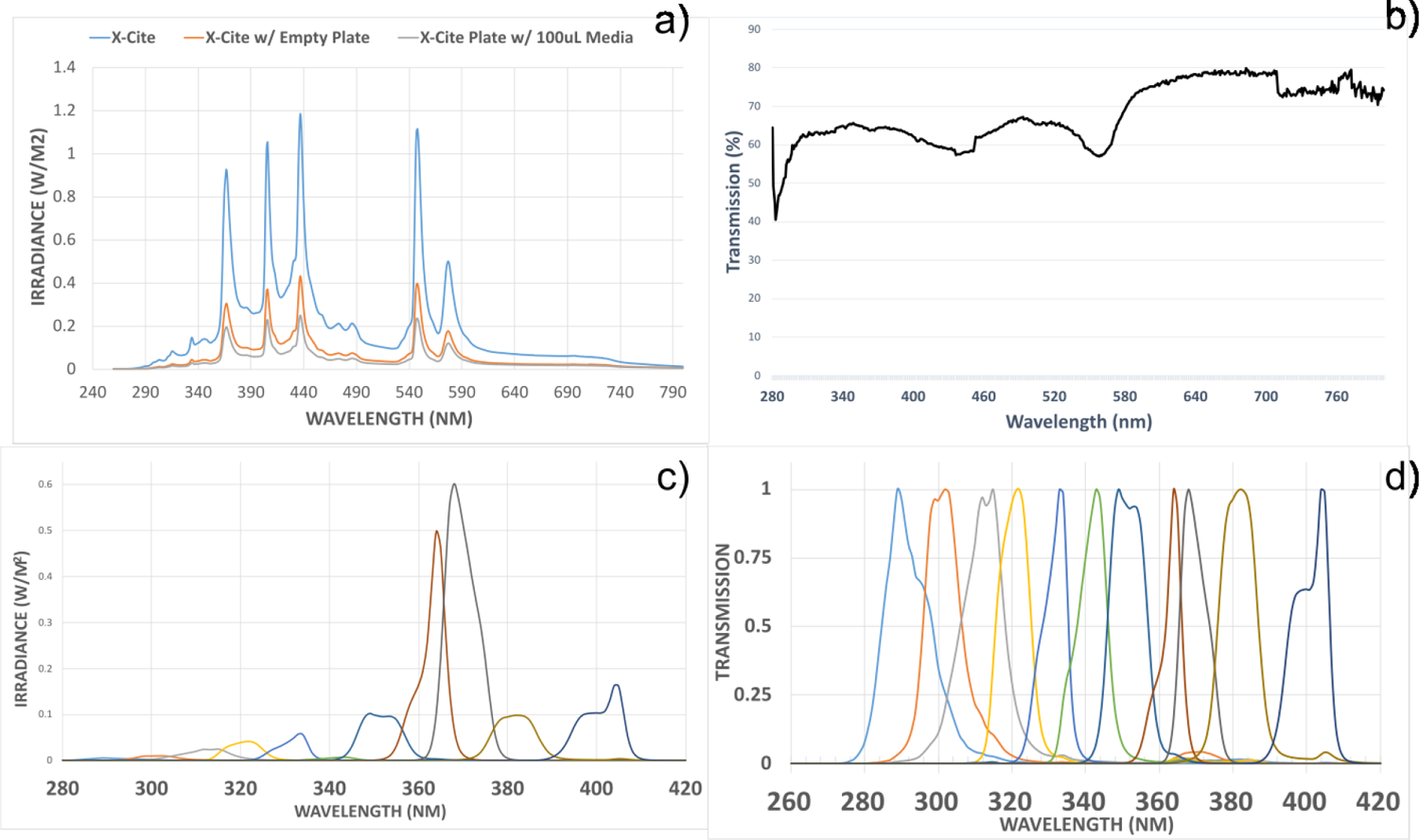
Spectral Characteristics of UVR Exposure Apparatus: a) light source spectrum when uninhibited as well as with cell culture plate and media in place. b) Spectral transmission profile of the media used in the study. c) Unadjusted irradiance data for the bandpass filters used in the study. d) Normalised transmission profiles of the bandpass filters utilised.

**Figure 7.**
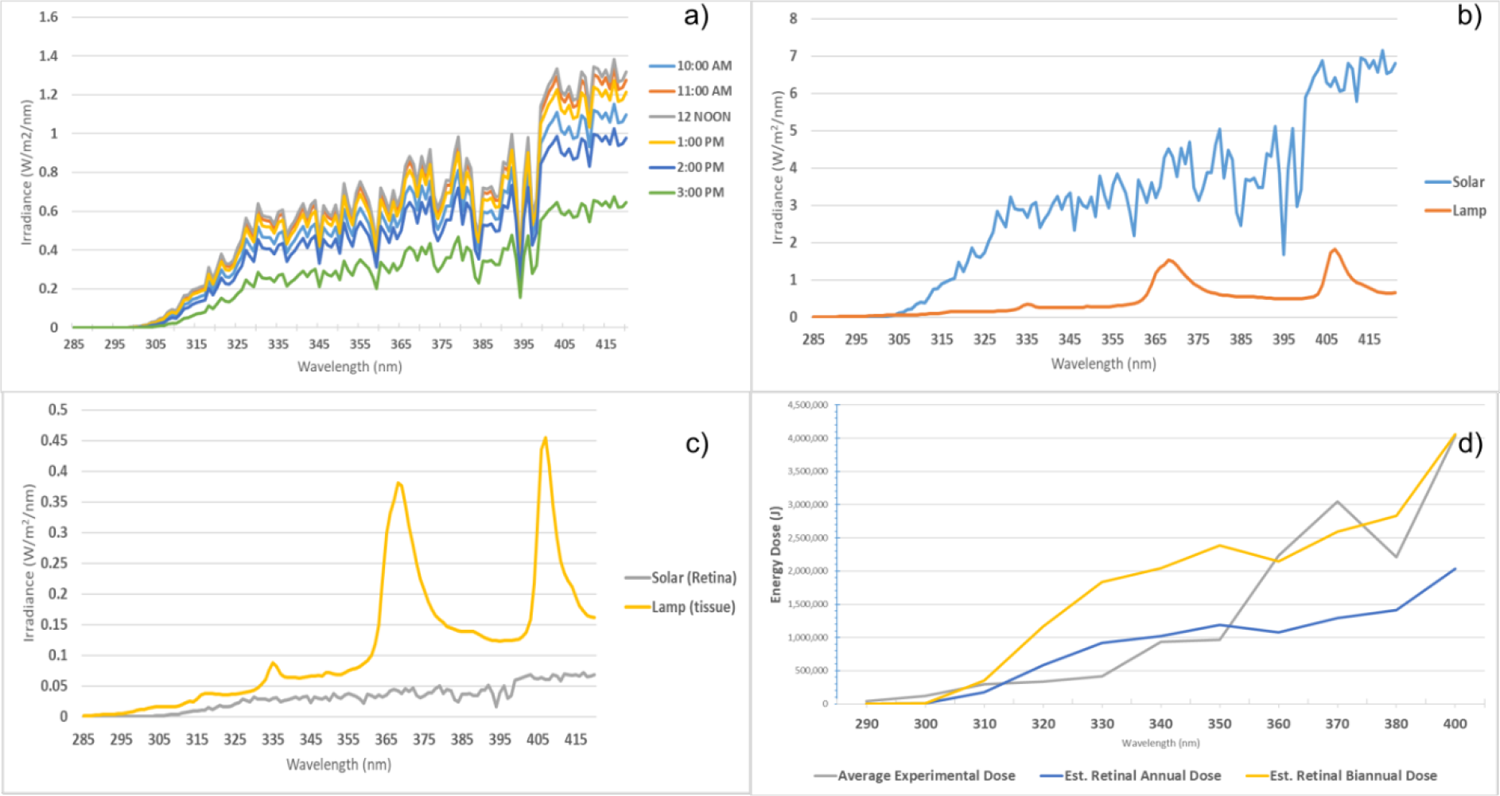
a) Estimated tropospheric ultraviolet radiation irradiance between 280 and 420nm at 0° N, 0° W, on 2020/06/21, between 1000-1500hrs. b) Total daily solar irradiance, from plot a) compared against metal-halide lamp output. c) Solar and lamp irradiances following transmission through the eye and culture plate, respectively. d) Average energy dose (grey line) of in vitro UVR exposures compared against annual (blue line) and biannual retinal dose (yellow dose).

The use of texture-based image processing techniques, such as the SER filter kernels, make it possible to reliably identify ridge or spot-like cell structures indicative of mitochondrial threads or punctate lysosomes in images with suboptimal signal-to-noise ratios. In the present study, by considering the ‘spot’ texture, it is possible to quantify pockets of ROS activity, condensed nuclear material and condensed mitochondria all of which are indicative of overt oxidative stress and pre-apoptotic cellular responses. When viewed in this way, the response spectra shows a concerted rise at 350 nm and 380 nm in all imaging parameters except the propidium iodide/Cell mask channel suggesting a sub-apoptotic response consistent with chronic oxidative stress (*Figure 5*; *plot d)*.

### RPE Response Falls along Distinct UVR Bands

To elucidate any possible high-dimensional clustering present in the data we undertook Monte-Carlo reference-based consensus clustering (M3C; John et al., 2020) to provide optimal identification of the number of clusters (represented by the letter *K*) within the data. Consensus clustering assumes that the optimal value for *K* is stable upon resampling, however, statistically robust methods to confirm stability have been lacking[12]. The M3C approach is founded upon the simulation of multiple reference null-data sets (i.e. *K*=1) based upon the real data supplied by the user and in so doing provides a null-hypothesis for significance testing when comparing higher values for *K*.

Using this method, based upon the coefficient data delivered by the linear models described previously, it was possible to simulate the cumulative distribution function (CDF) and proportion of ambiguous clustering (PAC) scores for the data. Finally, by comparing the real data against the simulated null-data sets, it is possible to generate a p-value regarding the probability of *K* = 1. The M3C analysis suggested that there are 3 or 4 stable clusters within the data, however, upon closer investigation it could be seen that one of the clusters identified referred to data of the blank control suggesting that the true number of clusters was 2 or 3.

Mapping the *K* = 2 clusters to the data simply divided the data into UV-Band UV-A bands. However, mapping clusters based on a *K*-value of 3 subdivided the UV-A response into 330-360 nm and 370-400 nm bands (*Figure 5*; *plot d*). This would suggest that the RPE’s UV-A sensitivity displays a distinct wavelength-dependent response which does not directly map to the established UV-A1 (340-400 nm) and UV-A2 (320-340 nm) bands, suggesting two possibly distinct mechanisms of action for each cluster.

## Discussion

In the present study we sought to describe the effect of UVR exposure upon the RPE by producing action spectra and response spectra relevant to parameters of interest within AMD pathology, specifically oxidative stress, cell viability and the tight junctional integrity of the RPE. Using the immortalised cell line, ARPE-19, we observed a previously unrealised sensitivity to UVR within the UV-A band, in particular at 350 nm and 380 nm.

The role of UVR in AMD remains a subject of some controversy, fuelled in part, by a lack of action spectra regarding the specific means by which UVR would precipitate AMD. Existing action spectra focus entirely on acute phase actions of UVR (i.e. fundus lesions) which are more relevant to occupational light hazards [18,19] than the chronic exposures expected from sunlight within a public exposure setting [20]. As a result, existing action spectra for the retina and the lens place little to no weight on UV-A effects, suggesting that only UV-B plays a role in UVR damage to these tissues[3,21].

Recently, Marie and colleagues generated action spectra between 390-520 nm for hydrogen peroxide (H_2_O_2_) and superoxide anion (O_2_^•-^) production in A2E-laden porcine RPE [6]. They were able to highlight that A2E acts as a potent photosensitiser capable of increasing H_2_O_2_ production 2-3-fold and doubling O_2_^•-^ content. A2E photosensitisation was highest at 415-455nm, closely following its absorbance spectrum [28,29]. This was complemented by a concomitant increase in expression of antioxidant genes such as *SOD2* and suppressed respiration rate as quantified via oxygraphy [6]. While A2E does have an absorbance peak within the UV-A band at 335nm, the researchers did not extend their action spectra beyond 390nm. However, it follows from the above results that a photosensitive effect would likely be observed if a broader spectrum were constructed.

Subsequently, Marie and colleagues investigated the spectral sensitivity of the neuronal retina in order to bring depth to our understanding of phototoxicity within retinal degeneration [30]. Through spectroradiometric investigation of the autofluorescent profile of each cell layer of the retina they demonstrated: 1) that the inner-segments of the photoreceptors are the most autofluorescent of the retinal cell layers, 2) that the mitochondria are the primary source of the autofluorescence and 3) that such autofluorescence is highest upon 350 nm excitation. The presence of defined peaks of autofluorescence and phototoxicity at 350 nm strongly imply the presence of a chromophore with maximal absorbance at, or within 10 nm of, 350 nm which resides within the retina. From their observation of peak autofluorescence at 350 nm Marie *et al*. reasoned that the putative chromophores behind the autofluorescence likely belong to the porphyrin or flavin families of photosensitisers [30]. Porphyrins normally exhibit a peak of absorbance around 400 nm, the Soret Band, with subsequent absorbance peaks found at longer wavelengths up to 630 nm, with little absorption within the UV spectrum. However, the molecule bacteriochlorin, a close relation of porphyrin, does exhibit dual absorption peaks at 349 nm and 372 nm [32]. Moreover, retinal accumulation of porphyrin-related compounds has been widely exploited in the use of photodynamic therapy to treat neovascular AMD, where the photosensitisers primarily occupy the choroid and RPE [33], thus providing a rationale for the role of porphyrin-related macromolecules in native retinal photosensitivity.

The observation of peaks in toxicity at 350 nm and 380 nm observed in the present study appear to be unique within the context of RPE phototoxicity. Previous studies, completed by Krohne *et al*., investigating the role of lipid peroxidation in autophagy, have shown RPE autofluorescent excitation maxima at 350 nm [39]. Moreover, the researchers demonstrated that the addition of photoreceptor outer segments (POS) increased excitation efficiency [39].

These findings would suggest that, if the dose was increased sufficiently, the cells would experience photodamage in a manner which reflects the excitation spectra and that the availability of lipids provided by POS would accelerate this process. Thus, the peak in effects observed at 350 nm in the present study could be a product of the same process of lipid peroxidation.

The peaks in cell death and mitochondrial dysfunction observed in the current study at 380 nm could point to a novel chromophore within the RPE. One candidate is the orphaned, UV-sensitive, circadian regulator OPN5 [42] which exhibits spectral sensitivity to 380 nm radiation. While expression of OPN5 has been confirmed in ARPE-19 [42] the foetal transcriptome of ARPE-19 [45] would suggest that OPN5’s role in the sensitivity of RPE to UVR may be limited to the very young. Further investigation, modulating OPN5 expression in RPE cell lines would prove useful in defining the role of OPN5 in long-wave UVR effects on the RPE.

While exposure to UVR is largely considered a hazard to be avoided, there remain some benefits to UVR providing the dose remains limited. Common examples of UVR’s therapeutic use include its role in vitamin D synthesis and its application as a therapy for infant jaundice. However, evidence of the benefits of UVR within the retina were, until recently, highly limited. Hallam et al. (2017) working with patient specific iPS-RPE derived from individuals harbouring a Y402H polymorphism within the *CFH* gene, demonstrated a possible benefit of long-wave UVR exposure to the RPE. In their study, patient-specific iPS-RPE cells and wild type were exposed to 4.5mW cm^-2^ of 390-410 nm radiation over the course of 5 days (for comparison, riboflavin/UVA corneal crosslinking uses approximately 3 mW cm^-2^ of 370 nm over 30 min [47]).Their results demonstrated a distinct difference in response to UVA/HEV exposure between the genotypes with the mutant line displaying an increase in expression of *SOD2, IL*6, *IL18 and IL1β* suggesting a pronounced inflammatory response. Moreover, imaging revealed that the mutant line showed significantly fewer vacuoles and significantly reduced drusen volume relative to the no-UVR control [46]. While it is currently unclear which chromophore or pathway is responsible for the beneficial responses this finding raises the possibility of using UVA/HEV radiation in a photobiomodulatory capacity to treat AMD within high-risk individuals.

The current study suggests that beyond the UV-B band, the wavelengths 350 nm and 380 nm are the most damaging to the ARPE-19 cell line in regard to cell viability and tight junctional integrity. As such, capturing discrete RPE sensitivity to UV-A vindicates the use of full UV spectral coverage when defining novel UVR effects upon the retina. Future work should focus on determining differential pathways of toxicity between these two wavelengths using mature retinal models and determine how such effects can be modulated through supplementation with small molecules relevant to AMD and oxidative stress.

## Materials and Methods

### Tissue Culture

The immortalised cell line ARPE-19 was purchased from the American Type Culture Collection (ATCC, MA, USA) and, following mycoplasma screening, expanded in T75 culture flasks using Dulbecco’s Modified Eagle Medium supplemented at a 1:1 ration with Ham’s F/12 solutions (DMEM F/12, Invitrogen, MA, USA) supplemented with 10% foetal calf serum (FCS, Merck, Germany). Once confluent, cells were passaged using porcine trypsin supplemented with ethylenediaminetetraacetic acid (EDTA, Life Technologies, CA, USA). For UVR exposure, ARPE-19 cells were seeded into black-walled (µClear, Greiner Bio One Ltd, Germany) and electrode embedded (96w20idf, Applied BioPhysics, NY, USA) 96-well plates at a density of 30,000 cells per well. Cells were maintained for at least 28 days before UVR exposure in DMEM with 4.5 g L^-1^ glucose, 1 mM sodium pyruvate and 1% FBS to allow for coherent cytoskeletal organisation and epithelial barrier formation[13,14].

### UVR Exposure

A full description of UVR exposure and calculations is given in the supplementary information/ Briefly, quantification of UVR at 1 nm intervals was carried out using an SR9910-v7 UV-VIS double monochromator spectroradiometer (Irradian Limited, Elphinstone, UK) fitted with a light guide and planar cosine corrected sensor assembly. The culture plate was irradiated inside a dedicated tissue culture incubator (Hera Cell, Heraeus, Germany), kept at 37°C, 5% CO_2_ and 100% humidity, with an uncollimated beam at 19 cm from the light-guide aperture of the UVR source, a 120 W mercury metal halide epifluorescence lamp (Excelitas Technologies Corp., NY, USA) Exposures took place over 72 h at three irradiance levels, one at full intensity (full irradiance) and two with ø21.3mm neutral density filters (ND 0.2 and ND 0.4 (ThorLabs Inc., NJ, USA)) mounted within a filter housing of the culture wells (*Figure 1*).

For each exposure, a negative (dark) control – comprised of a solid disc of black resin – was positioned in one of the available filter positions. Comparison of typical experimental UVR energy doses used in the present study and estimated terrestrial UVR dose confirmed that the UVR doses used in the present study achieved parity with the range of UVR doses experienced by a given person across the life-course.

### Cell Viability

Cell viability analysis was carried out using PrestoBlue™ (Invitrogen™, MA, USA) cell viability reagent according to the manufacturer’s instructions. In brief, culture medium was removed, and cells washed with phosphate buffered saline (PBS, Merck, MA, USA) containing calcium and magnesium chloride (referred to hereon as PBS^+/+^). Following washing, one well of the dark control cells was exposed to cell lysis buffer (RIPA Lysis and Extraction Buffer, Thermo Scientific, MA, USA) for 5 min at room temperature to act as a positive control. Next, 120 µL of 1:10 dilution of PrestoBlue™ (Invitrogen™, MA, USA) cell viability reagent and PBS^+/+^ was added to each exposed well. Cells were then incubated at 37°C and 5% CO_2_for 20 min to allow the assay to develop. Following incubation, 100 µL of the developed reagent was transferred to a solid white 96-well assay plate and fluorescence was read using a multi-modal plate reader (Ex 520nm/Em 580nm; GloMax Explorer, Promega, WI, USA).

### Electric Cell-Substrate Impedance Sensing (ECIS)

ARPE-19 cells were seeded onto ECIS culture-ware comprising 96 wells with each well housing 20 interdigitated 300 µm electrodes (96W20Eidf, Applied Biophysics, NJ, USA). Immediately prior to UVR exposure and following tissue maturation, the electrode plate was installed within a 96-well ECIS station. Spectral measurements of electrical impedance (Z), resistance (Ω) and capacitance (C) between 500 Hz and 64 kHz were determined at 11 min intervals during the UVR exposure.

### Cell Paint and Image Analysis

Within 2 h of UVR exposure, the cells were processed for imaging as follows: an equal volume of carboxy-H2DCFDA (Invitrogen™, MA, USA) solution diluted at a 1:500 ratio in Hank’s balanced salt solution (HBSS; Sigma-Aldrich, MI, USA) was added to the *in situ* culture medium and gently mixed by trituration to give a final carboxy-H2DCFDA dilution of 1:1000. The cells were then incubated with the staining solution for 30 min at 37°C. During incubation, a second staining solution comprising Hoechst 33342 (Sigma-Aldrich, MI, USA) diluted at a ratio of 1:500, CellMask™ Orange (Invitrogen™, MA, USA) diluted at a ratio of 1:500, MitoTracker™ Deep Red FM diluted at a ratio of 1:500 and propidium iodide (Sigma-Aldrich, MI, USA) diluted at a ratio of 1:1500 were made up in HBSS. During the last 10 min of the 30-min incubation, an equal volume of the second stain solution was added to the first solution in the wells. This was then incubated at 37°C for a further 10 min. When all staining was complete, 50% washes were performed in triplicate using HBSS.

Live imaging was performed using an automated microscope developed for high-throughput imaging (Operetta, PerkinElmer, MA, USA) the imaging chamber of which was maintained at 37°C and 5% CO_2_ throughout image capture. Seventeen fields were captured per well at 40X objective magnification (see *Figure 2* for representative images of fluorescent probes).

Image analysis was performed using the Columbus image analysis suite (PerkinElmer, MA, USA) Parameters of interest included number of cells, fluorescent intensity, texture analysis (saddle-edge-ridge (SER) textures) and STAR morphology (Symmetry properties, Threshold compactness, Axial properties, Radial properties, and profile) for each cell compartment. Following batch image quantification, all data were exported as a text file for analysis in Excel 2016 (Microsoft Systems, CA, USA) and R-studio[15].

### Data Curation and Statistical Analysis

PrestoBlue™ (Invitrogen™, MA, USA) data was scaled between the positive (lysed) and negative (no UVR) ‘dark’ controls using the formula: 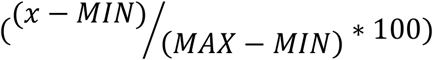 where ‘χ’ refers to the individual measurement, ‘MIN’ refers to the lowest value within the dataset and ‘MAX’ refers to the highest value. In a similar fashion, ECIS data were scaled between the positive (no-cell) and negative (no UVR) controls before being normalised to time zero (t_0_). Where possible, impedance, resistance and capacitance data were used to model Rb [tight-junction integrity], α [electrode coverage & cell adhesion] and Cm [membrane capacitance] values (collectively referred to as RbA) as described previously [16,17].

In order to model the UVR damage efficiency at each wavelength using the PrestoBlue™ (Invitrogen™, MA, USA) data, the processed cell viability for each wavelength and at each irradiance (full irradiance, ND02 and ND04) was used to calculate linear regression coefficients, the slope coefficient of which is indicative of the efficiency of UVR damage. Since the calculated slopes were all negative, they were first squared so they may be plotted logarithmically and normalised to the visible wavelength of 405nm.

Modelling photodamage efficiency using ECIS was performed by first choosing a ‘common action’ for all wavelengths as a 60% decrease in the electrostatic parameters (Z, Ω, C) from their original values. The time-point of the observations and the irradiance, enables the dose required to achieve the action to be calculated. The reciprocal of the dose required to fulfil the action (i.e. 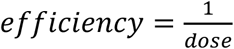)– at each wavelength provided an action spectrum for each of the electrostatic parameters. These action spectra were then normalised to 405nm to provide context in regard to visible radiation effects.

High content imaging data were averaged on a per wavelength basis, then normalised to the no UVR ‘dark’ control prior to action spectrum production. As outlined previously linear regression analysis was performed on the results from the three irradiance conditions (full irradiance, ND02, ND04) for each wavelength and the slope coefficient used to determine efficiency of response. Each exposure was replicated three times for each irradiance condition with four technical replicates present for each wavelength.

The data set was simplified using K-means clustering [32], supplemented by Monte-Carlo simulation driven consensus modelling, facilitated by the R package M3C[11], to provide empirical justification for the optimal value of *‘k’*.

All statistical analyses was carried out using R-Studio [15] and Excel 2016 (Microsoft systems, CA, USA), with data derived from three biological repeats each comprising four technical replicates.

## Supporting information

supplementary spectral data

