## supplementary spectral data for "Quantitative Action Spectroscopy Reveals ARPE-19 Sensitivity to Long-Wave Ultraviolet Radiation at 350 nm and 380 nm"

### Supplementary Data

#### Spectral Characterisation

Spectroradiometric quantification of UVR spectra was carried out using an SR9910-v7 UV-VIS spectroradiometer (Irradian Limited, Elphinstone, UK) fitted with a light guide and planar cosine corrected sensor assembly. The sensor was placed 19cm from the uncollimated light-guide aperture of the UVR source, a 120 W mercury metal halide epifluorescence lamp (Excelitas Technologies Corp., New York, USA) within a dedicated tissue culture incubator kept at 37°C, 5% $CO_2$ and 100% humidity.

Spectral scans between 260-800 nm, with 1 nm intervals, were performed to establish the source spectrum and the effects of the culture plate media on irradiance (**Error! Reference source not found.**;figure S1 plot a).

Based on the spectral readings, it was ascertained that at 19 cm from the sensor the light source achieved an irradiance of around 22 W $m^{-2}$ across the UVR region of interest (280-400 nm). In order to measure the influence of the culture plate on tissue irradiance, spectroradiometric readings were taken with the addition of a 96-well black-walled culture plate (#3603, Corning, New York, USA) placed within the illumination apparatus, maintaining a distance of 19 cm between the source and the spectroradiometer sensor, with the lid removed. This provided an irradiance of approximately 7 W $m^{-2}$..

Following the empty culture plate measurements, 100 µL of complete ARPE-19 culture media (DMEM/F12 (Gibco, MA, USA); 1% FCS (Sigma, MI, USA); 1%Pen/Strep (Invitrogen™, MA, USA) was added to each well of the plate and spectroradiometry repeated. This further reduced the irradiance to around 4.5 W $m^{-2}$. Based on these data, the transmission of the media in the

280-400 nm range was calculated to be 60-65% (**Error! Reference source not found.**, *panel d))*. By subtracting the difference in irradiance between the empty plate, and the plate with media added, from the unimpeded source irradiance it is possible to approximate the irradiance reaching the surface of the cultured cells to be 19.4 W $m^{-2}$.

### 3D Design & Manufacture of Bandpass Filter Housing

Computer aided design (CAD) software (Inventor 2018, Autodesk Inc., CA, USA) was utilised to design a housing that would allow the Ø21.3mm bandpass filters to be placed under a standard 96-well black-walled culture plate such that cultured cells could be irradiated from below.

The design of the filter holder ensured that each of twelve filter positions covered four wells of a standard 96-well culture plate, thus providing technical replicates for each exposure (**Error! Reference source not found.**; panel b). Moreover, in order to protect the filters from the humidity within the incubator, two quartz windows were installed on either side of the filter housing within the holder (see**Error! Reference source not found.**; plot b for quartz transmission data). To add further protection, the filters were stored within a desiccant container between exposures to extract any moisture caught within the cavities of the filter.

### Calculating Tropospheric UVR

The TUV model used in this study *(TROPOSPHERIC ULTRAVIOLET VISIBLE (TUV) MODEL (version 5.0) S. Madronich et al., Atmospheric Chemistry Division National Centre for Atmospheric Research P. O. Box 3000, Boulder, Colorado)* was downloaded from the NCAR website *[https://www2.acom.ucar.edu/modeling/tropospheric-ultraviolet-and-visible-tuv-radiation-model]* and installed on a computer running Windows 10. All atmospheric inputs were left at their default settings, longitude and latitude were kept at 0° and the model date set to 21/06/2020.

### Calculating UVR Irradiance of the Retina

Based upon published evidence [4,5] we selected a working irradiance of between 0.1-1% of the UVR incident upon the eye reaching the retina. Moreover, based on these findings and the measured transmission properties of the culture media and the black wall plate it is possible to directly compare ambient UVR with the *in vitro* irradiance utilised in the present study.

Upon comparison it can be seen that the estimated *in vitro* tissue irradiance was consistently higher than the estimated solar UVR irradiance of the retina *in vivo*, the difference is especially large around the mercury emission peaks at 370 and 405nm where the *in vitro* irradiance is approximately 4.5-fold that of the estimated retinal UVR irradiance (**Error! Reference source not found.Error! Reference source not found.**; plot c). However, the example TUV solar irradiances presented here are instantaneous values at five time-intervals around midday (**Error! Reference source not found.**; panel a). When considering the uninhibited cumulative dose over two years compared to the dose achieved in exposures used throughout the project the output of the two sources begin to converge (**Error! Reference source not found.Error! Reference source not found.**; plot d).
